# Tracking trends in monarch abundances over the 20^th^ century is currently impossible using museum records: a response to Boyle et al. (2019)

**DOI:** 10.1101/581801

**Authors:** Leslie Ries, Elise F. Zipkin, Rob P. Guralnick

## Abstract

The onslaught of opportunistic data offers new opportunities to examine biodiversity patterns at large scales. However, the techniques for tracking abundance trends with such data are new and require careful consideration to ensure that variations in sampling effort do not lead to biased estimates. The analysis by Boyle et al. (2019) showing a mid-century increase in monarch abundance followed by a decrease starting in the 1960s used an inappropriate correction with respect to three dimensions of sampling effort: taxonomy, place, and time. When the data presentenced by Boyle et al. (2019) are corrected to account for biases in the collection process, the results of their analyses do not hold. The paucity of data that remain after accounting for spatial and temporal biases suggests that analyses of monarch trends back to the beginning of the 20^th^ are currently not possible. Continued digitization of museum records is needed to provide a firm data basis to estimate population trends.

---

Opportunistic data provide a tantalizing opportunity to examine patterns in biodiversity over large spatiotemporal scales (Soroye et al. 2018). Recent methodological advancements hold promise for utilizing such data to estimate trends while also highlighting the difficulty in accurately assessing biases (e.g., Bartomeus et al. 2013, Kramer-Schadt et al. 2013, Dorazio 2014). The idea is to determine the total number of collections of similar species within a reasonably comparable time and place to correct for variations in sampling effort. Using specimen data collected opportunistically, Boyle et al. (2019) showed a mid-century increase in monarch abundance followed by a decrease starting in the 1960s. However, their analysis used an inappropriate correction with respect to three dimensions of effort: taxonomy, place, and time. When these data are re-standardized to account for biases in the collection process (Fig. 1), the pattern presented disappears entirely (Fig. 2).

**Figure 1.**
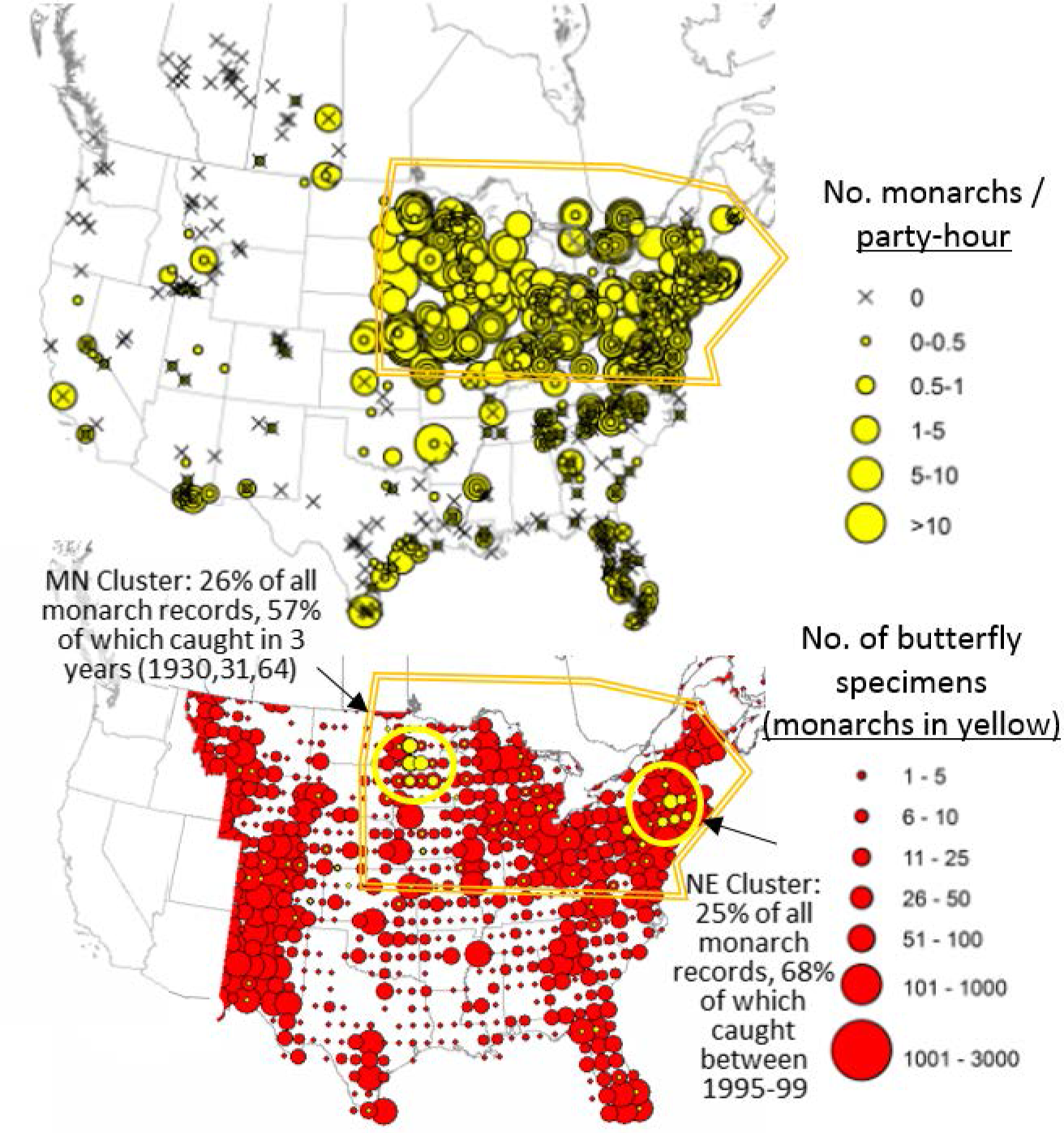
(a) Within the US and Canada, monarchs are most abundant during the summer in the northern part of their range (outlined in orange) as shown by a corrected abundance index from an annual nationwide survey by the North American Butterfly Association (2003-2017). (b) The number of records from museum collections is shown throughout the range used in Boyle et al. (2019), where they divided the total number of monarch specimens (yellow circles) by the total number of specimen records for all Lepidoptera (red circles show records for all butterflies). Note that 51% of all monarch records were collected in two spatially restricted clusters, and within those clusters, the majority of records are from a restricted set of years, leading to extreme spatial and temporal bias in the collection effort.

**Figure 2.**
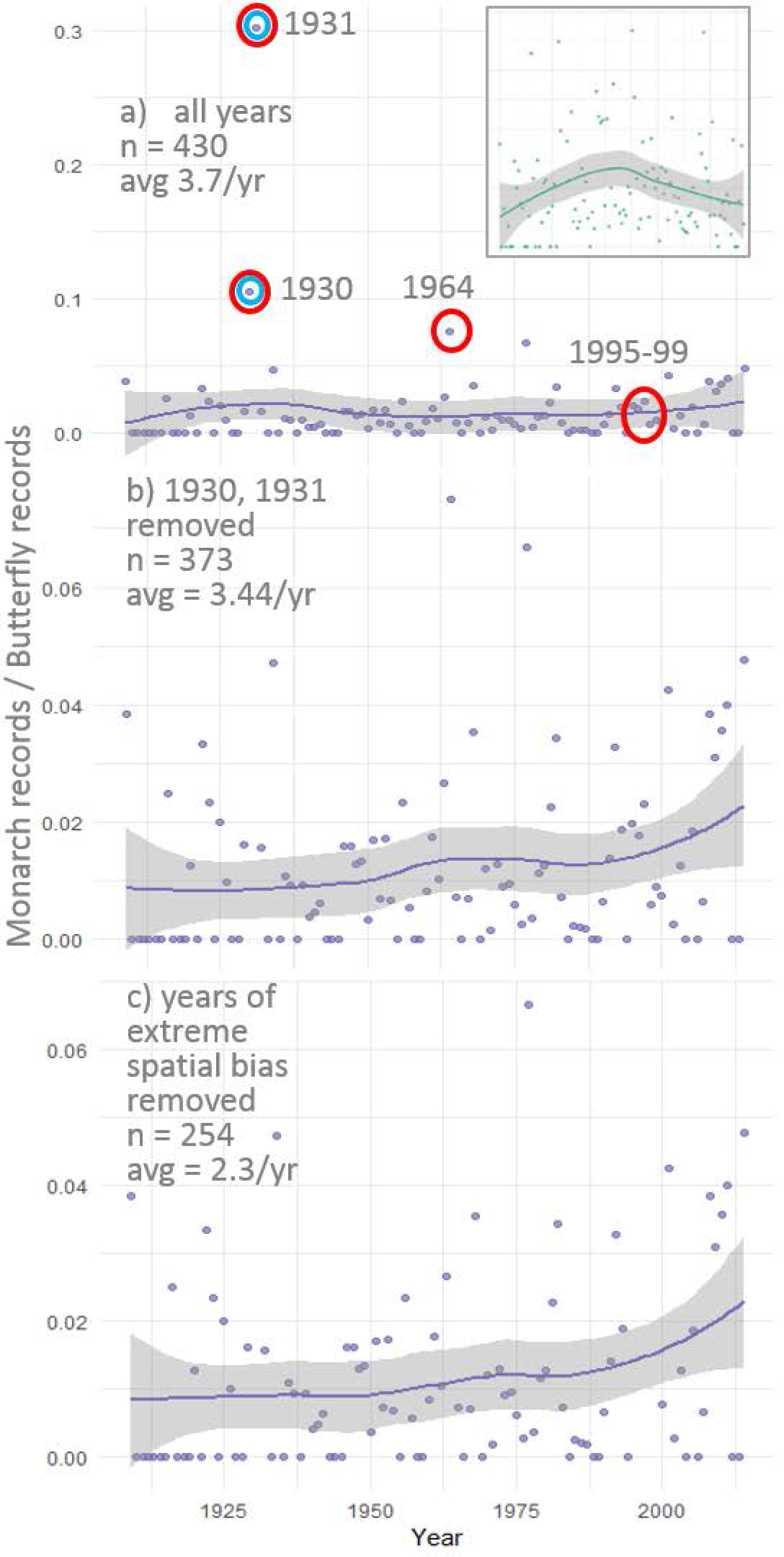
The number of monarch observations corrected by all butterfly records during the same year. Data were restricted to primary monarch breeding grounds (east of −100W, between 38 and 50N, outlined in orange in Fig. 1) during the summer when they are most abundant (between June 14^th^ and August 15^th^). Original pattern reported in Boyle et al. (2019) shown in inset, reprinted from Wepprich (*in review*) with permission. All data came from Boyle et al. (2019) and were fit using the same default LOESS smoother with 95% confidence intervals in the *ggplot2* R package (Wickham 2016). (a) Results when all years included. Note that we highlight years removed from the Boyle et al. (2019) analysis (blue circles) and years in which the major collection efforts occurred in the Minnesota cluster (1930,1931, 1964) and the North East cluster (1995-1999) (red circles; see also panel b). (b) Results from an analysis in which we remove outlier years as defined by Boyle et al. (2019; circled in blue in panel a). (c) Results from an analysis in which we remove both outlier years defined by Boyle et al. (2019) and years with extreme spatial bias in collection efforts (circled in red in panel a). The total number of monarchs included in each analysis and average number per year is shown in the top left corner of each panel.

Moth records should generally not be used as a correction effort for butterfly species because the collection process for moths (light trapping at night) is substantially different than for butterflies (net captures during the day). Wepprich (*in review*) demonstrates that the pattern presented in Boyle et al. (2019) could be explained by a spike in moth museum records in the 1950s. Here, we additionally address the issue of spatiotemporal bias in the collection process. One way to reduce this bias is to restrict analysis to the core range when the species is most evenly distributed and consistently abundant. Records falling outside core ranges do not add significant information but risk collection biases, which can lead to spurious patterns. The eastern population of the migratory monarch reaches their highest abundance and most consistent distribution during the summer (Fig 1a) and constraining abundance estimates to this range has been shown to best capture trends (Saunders et al. 2019, Ries et al. 2015, Inamine et al. 2016).

Using the same dataset as Boyle et al. (2019), we excluded moths and then examined patterns in monarch collection locations compared to all butterflies (Fig. 1b). Even after confining the analysis to their core range (orange boundary, Fig. 1), we found overwhelming spatial and temporal bias in the specimens. The majority of all summer monarch records (51%) were accumulated in two restricted clusters in Minnesota and New England with most specimens collected during a very limited number of years (Fig. 1b). We find no mid-20^th^ century spike in abundance when using the corrected data (Fig. 2c). Note that with appropriate correction, the average number of observations per year is only 2-3 monarchs (Fig. 2c).

The spurious patterns in the monarch dataset represents a cautionary tale in the use of opportunistic data to track population trends. Based on current, limited data availability (Fig. 1b, 2), using digitized museum records to track the monarch butterfly populations over the last century is not possible due to overwhelming spatiotemporal bias (Fig. 1b, 2). We strongly support continued efforts to digitize monarch specimens which may eventually provide the proper basis to explore long-term monarch population trends.

